# Network pharmacology-based discovery and experimental validation of novel drug repurposing candidates in Alzheimer’s Disease

**DOI:** 10.64898/2026.03.05.709917

**Authors:** Attila Jones, Tina Loeffler, Evan Wu, Vijay R. Varma, Hae Kyung Im, Madhav Thambisetty

## Abstract

Despite a growing body of evidence implicating genetic variants and proteins encoded by them with risk and pathogenesis of Alzheimer’s disease (AD), this knowledge has not been successfully translated into effective AD treatments. We integrated current genomic, transcriptomic and proteomic profiles of AD into a network pharmacology framework that leverages comprehensive gene-gene and drug-target interactions. This approach allowed us to screen 2,413 drugs for repurposing opportunities in AD. Computational validation and drug prioritization was followed by experimental validation in 33 cell culture-based phenotypic assays combined with Bayesian hypothesis testing. Our network-based screen rediscovered drugs in clinical trials for AD, providing computational validation. Besides many cancer drugs, the screen identified three drugs previously implicated in AD-related endophenotypes: the primary bile acid chenodiol, arundine (3,3’-diindolylmethane), and cysteamine. In analysis of results from culture-based phenotypic assays, large Bayes factors supported the hypothesized benefits of arundine and the chenodiol derivative, tauroursodeoxycholic acid (TUDCA), in amyloid-***β*** clearance and release and neuroinflammation. Follow-up network analyses mechanistically implicated Regulator of G protein signaling 4 (RGS4) in the plausible therapeutic actions of arundine and TUDCA. A network pharmacology approach identified TUDCA and arundine as promising repurposing candidates in AD that rescue disease-relevant molecular phenotypes by acting on AD-associated genes through regulation of G protein signaling.

## 1 Introduction

Alzheimer’s disease (AD) is a progressive neurodegenerative disorder and the most common form of dementia affecting a growing number of individuals in ageing societies for which effective treatments are unavailable [1, 2]. Repurposing drugs approved for other indications is a promising approach towards identifying effective disease-modifying AD treatments [3–5]. One of the milestones of the National Plan for Alzheimer’s Disease is to “Initiate research programs for translational bioinformatics and network pharmacology to support rational drug repositioning” [6].

Although a few major genetic risk factors for AD are known—most importantly the *ϵ*4 allele of the apolipoprotein E gene (APOE4) [7]—our understanding of the early etiological triggers of the disease is still limited. Among several reasons for this lack of clarity are the considerable genetic and cellular complexity of AD, and the confounding of early etiological processes by compensatory and degenerative pathologies over the decades-long disease progression [8]. Consequently, discovering treatments targeting early causal drivers of AD pathogenesis has been a major hurdle to developing effective treatments. In this study, we attempted to address this challenge in two ways. First, we derived plausible mechanistic insights into AD from previous genome-, transcriptome- and proteome-wide association studies (GWAS, TWAS, PWAS), comparing AD and control samples, including analyses of differentially expressed genes (DEGs) in APOE3/3 versus APOE4/4 individuals, as well as comparison of DEGs in iPSC-derived neurons, astrocytes, and microglia from APOE3/3 versus APOE4/4 individuals [5, 9–16] (Fig. 1A).

**Fig. 1.**
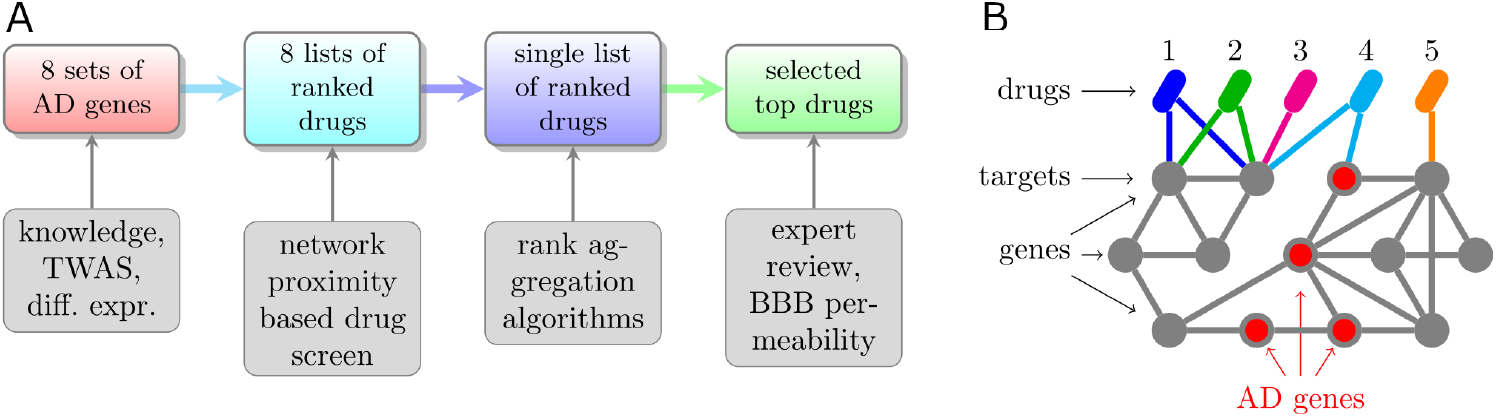
(A) Workflow of computational drug repurposing in Alzheimer’s disease (AD). (B) Principle of network proximity-based drug screen for AD. Drug 4 and 5 are plausible candidate AD drugs because they target either directly or indirectly an AD risk gene (i.e a gene mechanistically involved in AD), respectively. On the other hand, drugs 1–3 only target genes at least three interactions away from AD genes and therefore are likely not plausible candidate AD drugs.

Second, we exploited rich mechanistic information represented by two interconnected networks (Fig. 1B): (1) the network formed by all drugs and their target genes (drug-target network), and (2) the human gene-gene interaction network—known as the interactome—whose AD subnetwork (AD module) was defined from AD risk genes derived from the above omic studies [17]. Combining these two networks, we estimated the potential efficacy of each drug for AD by evaluating its network proximity [18], which is based on the drug’s network topological distance from the AD module. Network proximity has been shown to be a powerful systems-level alternative to purely correlative approaches to computational drug repurposing [19]. Using manual prioritization of top-scoring drugs, we complemented this computational screen with experimental validation. Our results identify the primary bile acid chenodiol, organosulfur compound cysteamine, and the indole-3-carbinol metabolite arundine as promising AD treatments.

## 2 Methods

### 2.1 AD risk gene sets

Given an AD risk gene set, the network structure of the human interactome, and another network formed by drugs and their targets (drug-target network), the topological proximity of a given drug to the AD risk gene set can be quantified [18] and used as evidence for repurposing that drug for AD. Rather than compiling a single AD risk gene set from multiple studies, we evaluated drug proximity separately for multiple AD risk gene sets and then aggregated the multiple proximity values (see Discussion). Table S1 summarizes the AD risk gene sets used in this study. The genes in each set are listed in Data file 1.

### 2.2 Previous and novel TWAS on AD

See Supplemental Methods.

### 2.3 Drug-target network

We queried ChEMBL [20] (v29, July 2021) for phase 3 and 4 drugs and their targets that satisfied the following conditions: (1) the target is a human protein and (2) its average *−* log_10_ Activity or *p*Activity is ≥ 5, which corresponds to ≤ 10 *µ*M following [19]. Average *p*Activity is defined as 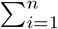 *p*Activity_*i*_*/n*, where *n* is the number of activity measurements for a given drug-protein pair and Activity_*i*_ is the standard value of the *i*-th measurement stored in the `activities.standard_value` SQL variable.

The result of these operations is a set of 25,329 drug-target pairs, which constitute the drug-target network.

### 2.4 Gene-gene interaction network

We used the network compiled by [21], which consists of 11,133 nodes (genes) and 217,160 edges (gene-gene interactions).

### 2.5 Network proximity calculations

[18] developed the network proximity method for in silico drug efficacy screening. It quantifies the topological proximity *d* in the interactome of the set *T* of a drug’s target genes to the set *S* of disease genes:

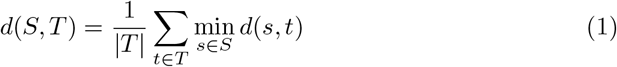

where *d*(*s, t*) is the shortest path length between disease gene *s* and drug target *t*. The equation can be interpreted as the average length of shortest paths taken from all targets of a drug to all the disease genes.

We used [our modified clone](https://github.com/attilagk/guney_code) of the original, now obsolete, implementation of [18], which allows running the code under Python 3.

### 2.6 Rediscovery rate of drugs for a disease indication

Let *T*_*k*_ be the set of top-*k* ranked drugs and *B*_*l*_ that of the bottom-*l* ranked drugs. We define 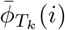, the average clinical phase of the top-*k* drugs for disease indication *i*, as follows:

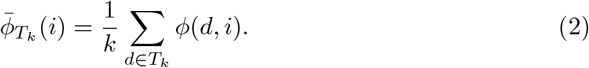

Moreover, we define 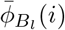, the average clinical phase of the bottom-*l* drugs for *i*, analogously. Given disease indication *i*, we then define the rediscovery rate of the top-*k* drugs relative to the bottom-*l* drugs as:

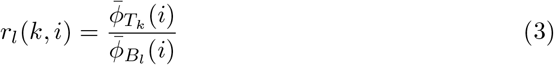

The notation *r*_*l*_(*k, i*) emphasizes that rediscovery rate depends on disease indication *i* and on the top-*k*/bottom-*l* choices. In Fig. S6, we held *l* = 1808 constant and varied *k*.

### 2.7 Rank aggregation

As Fig. 1 shows, we aggregated eight lists of network-proximity-ranked drugs into a single list with rank aggregation algorithms. We performed rank aggregation with the TopKLists R package several times, each time with a different algorithm. We then assessed algorithm performance with modified Kendall distance and found MC3 to perform best (Fig. S5). Finally, we used the MC3-aggregated list in our workflow.

### 2.8 Cell-based assays

We performed dose-response analysis to test whether selected drugs can rescue molecular phenotypes relevant to AD in 33 cell-based assays grouped into nine experiments [22, 23]; see Supplemental Methods.

To evaluate chenodiol, we used TUDCA, the taurine-conjugated epimer of chenodiol, because prior studies suggest neuroprotective effects in neurodegenerative disease models [24] and in clinical trials for ALS [25]. To evaluate cysteamine, we used cysteamine hydrochloride, which is FDA-approved for corneal cystine crystal deposits in patients with cystinosis [26, 27] and has improved renal function in pediatric cystinosis [28]. For arundine, we used the parent compound [29, 30].

In dose-response analyses, each drug was applied in three concentrations (in *µ*M): TUDCA 1, 10, 100; cysteamine-HCl 1, 3, 10; arundine 1, 3, 10.

Table S4 lists all 33 assays and, for each assay, whether the desired neuroprotective effect corresponds to increased or decreased bioactivity. In all assays, drug-treated cells were compared with either vehicle control (VC) or lesion control.

For each drug-assay pair, we evaluated posterior probability *P* (*H*_*i*_|data) of protective, neutral, and adverse effect (formal hypotheses *H*_1_, *H*_0_, *H*_2_). We modeled dose-response relationships (*x*-*y* data) with the Bayesian nonlinear regression model shown in Fig. S8:

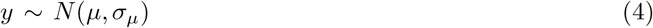

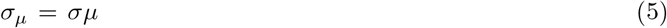

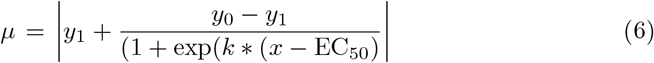

We interpret parameters *y*_0_ and *y*_1_ as expected bioactivity at zero and saturating concentration and define fold change as FC_*y*_ = *y*_1_*/y*_0_. We used a gamma prior density for fold change:

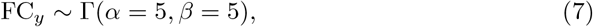

and performed Bayesian estimation with the default Markov chain Monte Carlo sampler settings in PyMC (https://www.pymc.io/). For efficient sampling, each bioactivity dataset was standardized so that 10 units corresponded to one standard deviation. We inspected posterior fitted curves, effective sample sizes, 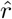, and Markov chain standard errors, and excluded three poorly fitted datasets out of 91 total datasets. Posterior fold-change densities are shown in Fig. S9.

We defined hypotheses *H*_1_, *H*_0_, *H*_2_ and used posterior fold-change density to calculate posterior probabilities in the Supplementary Methods. Eq. 7 implies prior mean fold change of 1 (no effect). Hypotheses were defined so that each drug has prior probability 0.2 of protective or adverse effect and 0.6 of neutral effect.

From posterior probabilities, we calculated the posterior odds (Bayes factor):

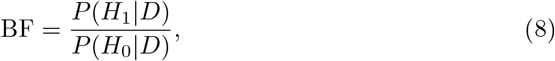

where *D* denotes dose-response data. We used twice the log-scaled Bayes factor, 2 *×* log BF, to quantify evidence for a protective effect.

## 3 Results

### 3.1 Identifying drugs with the strongest potential efficacy for AD

Drugs proximal to a disease module in the human interactome are good candidates for repurposing to that disease [19]. We performed a network-proximity-based repurposing screen for AD using 2,413 ChEMBL drugs that were either approved by the US Food and Drug Administration or tested in phase 3 clinical trials for any indication (Fig. 1A).

We evaluated each drug’s network proximity to each of eight AD modules (or AD gene sets), derived from one curated knowledge-based source and seven multi-omic sources (Table S1). For each AD module, we obtained a proximity score vector and ranking of all 2,413 drugs, where drugs with strongest potential efficacy for AD were at or near rank 1 (Fig. 1A, Fig. 2A-B, Data file 2).

**Fig. 2.**
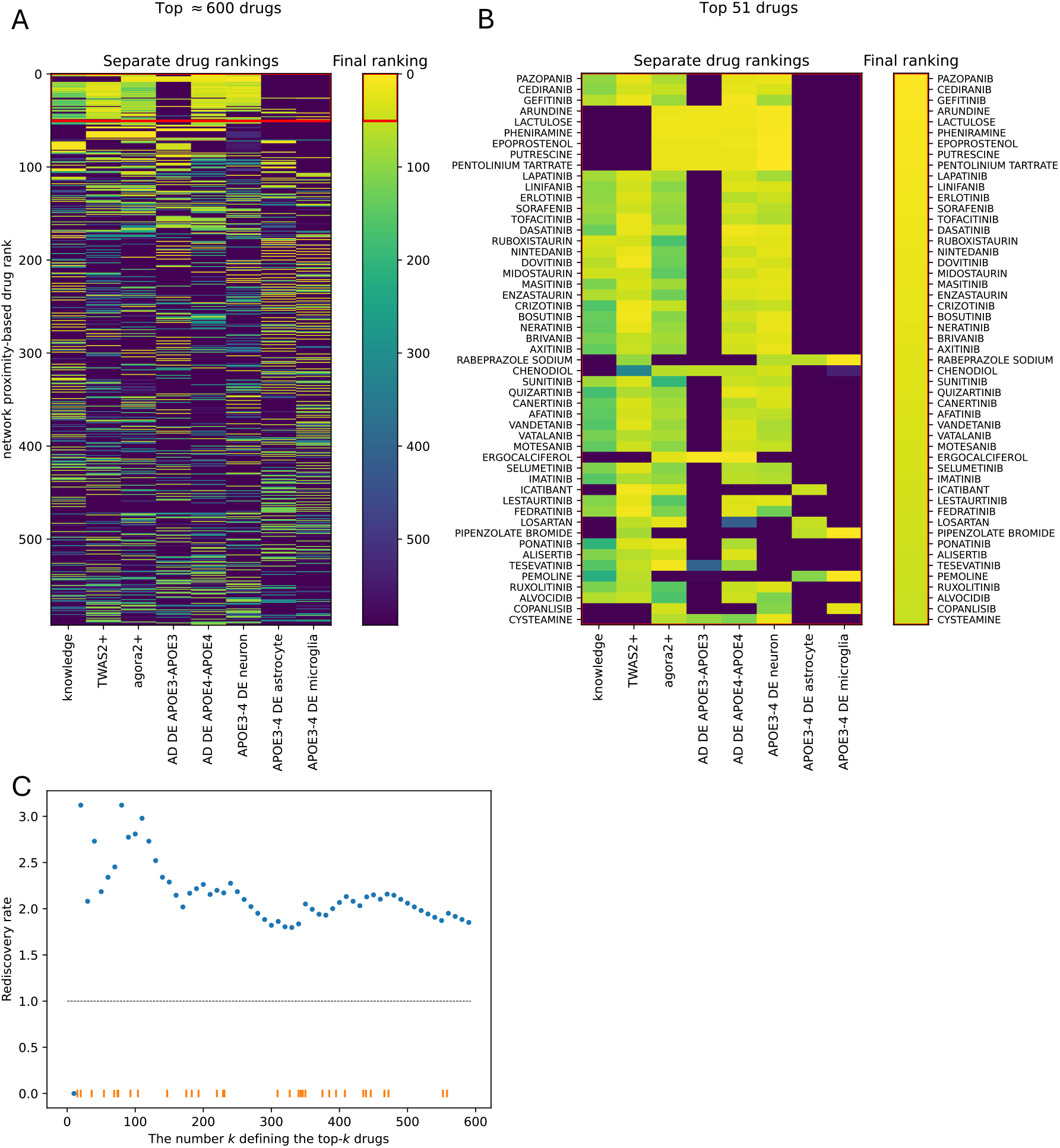
Computational drug screen and computational validation. (A-B) Scoring and ranking 2413 drugs based on their network proximity to various AD risk gene sets. Eight AD risk gene sets were input to the workflow. The separate drug rankings across imput AD risk gene sets were aggregated into a single final ranking, which is color coded (yellow: top-ranked drugs, blue: bottom-ranked drugs). (A) shows the top ≈ 600 drugs while (B) only the top 51. (C) Rediscovery rate *>* 1 for the top-*k* drugs suggests these drugs, highly ranked by network proximity, are enriched in drugs already investigated or approved for AD.

Although the eight input AD risk gene sets had relatively few shared genes (Fig. S3), the corresponding network-proximity score vectors were highly correlated (Fig. S4). Consequently, the eight drug ranking lists were substantially similar (Fig. 2A). This suggests that even with limited overlap in AD genes, corresponding network modules are sufficiently close in topological space to yield similar drug proximity patterns.

Assuming equal relevance of all eight AD gene sets, we aggregated the corresponding rankings with uniform weights. This yielded a final ranked list of the top 605 drugs (Fig. 2A, Data file 2). Top drugs in the final list also tended to rank highly in a majority (five of eight) of gene-set-specific rankings (Fig. 2B, Fig. S1).

### 3.2 Computational validation of network proximity-based drug screening

To validate our network-based drug screen, we tested whether highly ranked candidate AD drugs were enriched among drugs already investigated in or approved for AD. We quantified enrichment with rediscovery rate (Methods, Eq. 3), comparing drugs in top-*k* versus bottom 1,000 ranks. Fig. 2C shows rediscovery rates for AD of roughly twofold to threefold in top-ranked drugs (depending on *k*), indicating that the network-proximity approach recovers drugs with biologically plausible AD relevance while still nominating novel candidates.

We next assessed rediscovery for 35 non-AD indications with sufficient data (at least 100 drugs approved or in phase ≥ 1 clinical trials; Fig. S6). Several indications showed enrichment patterns similar to AD, especially cancers, while neuropsychiatric indications such as major depressive disorder, schizophrenia, and Parkinson’s disease did not. These findings suggest that candidate AD drugs from this network-based screen may target molecular mechanisms that overlap with diseases such as cancer.

### 3.3 Manual prioritization of top-ranking candidate AD drugs

We manually prioritized top-scoring drugs based on (1) blood-brain barrier permeability from the BBB database [31] and (2) prior evidence for roles in AD-relevant mechanisms (Table S2). This yielded three lead candidates: chenodiol, arundine, and cysteamine.

### 3.4 Experimental validation of candidate AD drugs

We evaluated whether chenodiol, arundine, and cysteamine rescue AD-relevant molecular outcomes in 33 cell culture-based phenotypic assays (Fig. 3). In these studies, we evaluated chenodiol as TUDCA (its taurine-conjugated epimer), based on prior evidence suggesting neuroprotective effects in neurodegenerative disease models (Methods).

**Fig. 3.**
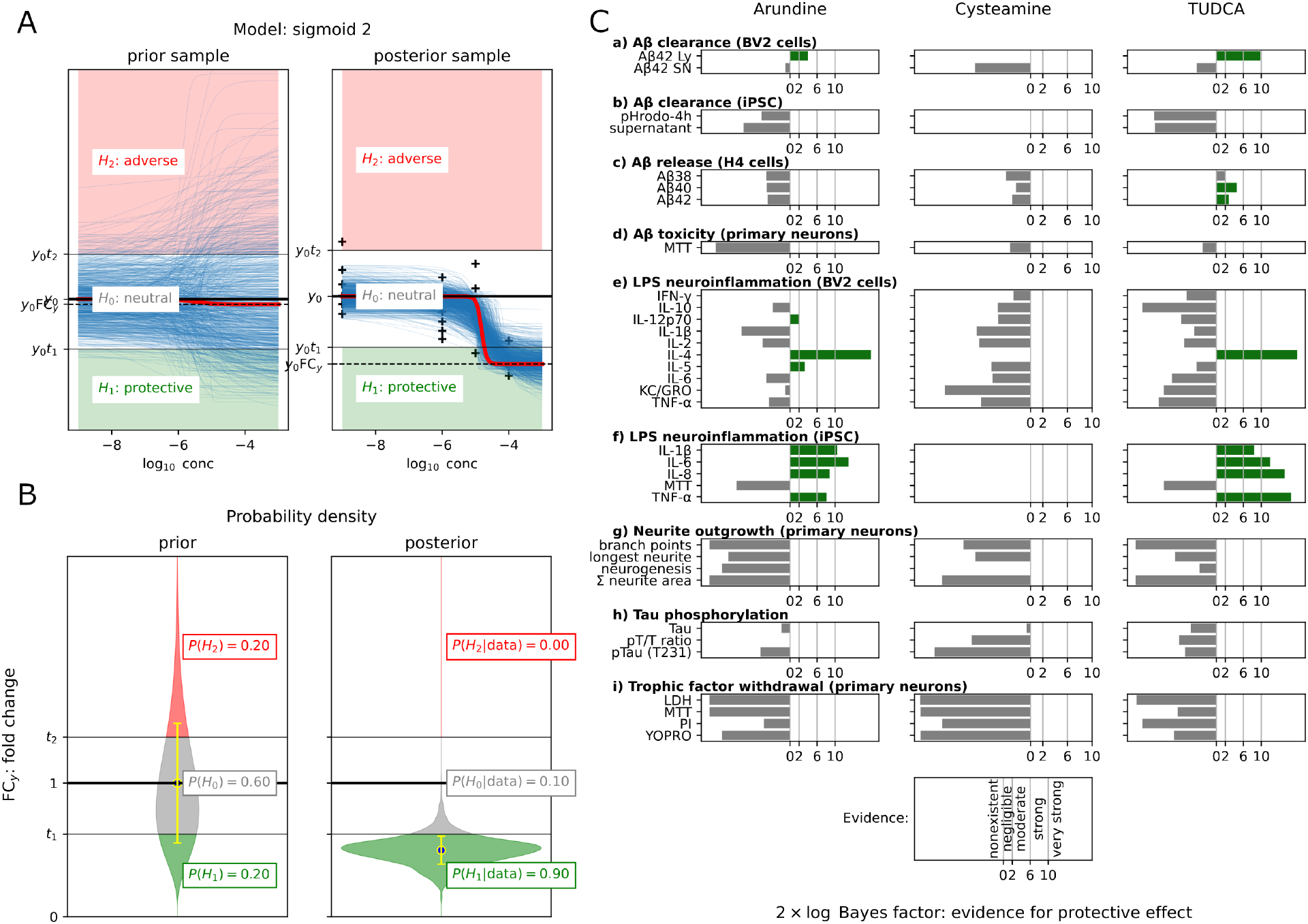
Experimental validation. Assessing the hypothetical neuroprotective effect of Arundine, Cysteamine and TUDCA in 33 cell-based assays from 9 experiments, labeled as (a)–(i). Note that TUDCA is the taurine conjugated epimer of Chenodiol. Green bars indicate moderate to very strong evidence for protective effect as quantified by the Bayes factor, which is the posterior odds of protective and neutral effect. Blank (white) slots indicate either that the experiment was not performed on the drug or that model fit was poor.

For each drug-assay pair, we analyzed dose-response data (Methods, section 2.8; Fig. S9, Fig. S10) and quantified evidence for protective effects with twice the log-scaled Bayes factor, 2 *×* log BF (Fig. 3).

We observed moderate to very strong evidence that arundine and TUDCA, respectively, enhance A*β*1-42 clearance in BV2 microglial cells (Fig. 3a). We found moderate evidence that TUDCA reduces release of A*β*40 and A*β*42 in H4-hAPP cells (Fig. 3c). We observed moderately strong evidence that arundine protects BV2 cells from LPS-induced neuroinflammation (Fig. 3e). We also found strong to very strong evidence that both arundine and TUDCA lower LPS-induced neuroinflammation in iPSC-derived adult human AD microglia (Fig. 3f).

## 4 Discussion

In this work, we report that TUDCA, the taurine-conjugated epimer of chenodiol, and arundine are promising repurposing candidates in AD. This conclusion is supported by network-proximity analysis and experimental validation in multiple AD-relevant phenotypic assays.

As shown in Fig. 4 and Table S3, network paths between prioritized drug candidates (chenodiol, arundine, cysteamine) and AD genes reveal Regulator of G protein signaling 4 (RGS4) as a key target linking these drugs to AD. RGS4 itself may be considered AD-relevant because it is differentially expressed in AD brains [5, 32] and has been implicated in beta-site amyloid precursor protein processing pathways.

**Fig. 4.**
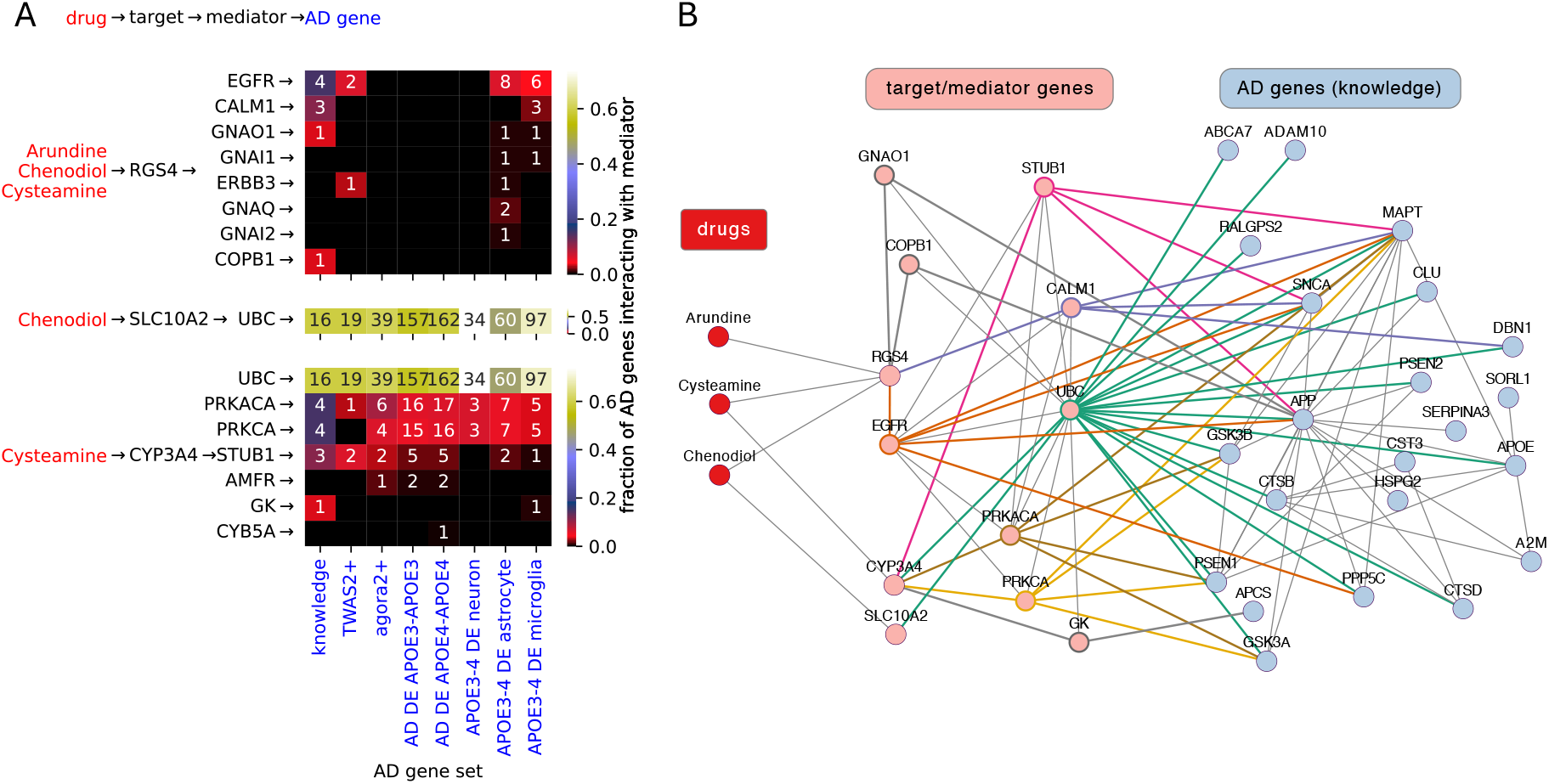
RGS4 (Regulator of G protein signaling 4) is a key target of Arundine, Chenodiol, and Cysteamine, suggested by network paths between these drugs and AD genes. The figure depicts only three edge-long paths, which all have the form: drug 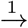 target 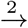 mediator 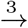 AD gene. Such paths are both relatively frequent and short to explain potential pharmacological effects on AD genes. There are only 4 two edge-long paths; all of those involve RGS4 as target (drug 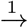 RGS4 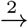 AD gene, see Table S3). Moreover, RGS4 can itself be considered an AD gene based on its membership in two of the eight AD gene sets of this study (one edge-long paths, Table S3). (A) The top, middle and bottom panel shows mediators (rows, *y* axis) interacting with drug targets RGS4, SLC10A2, CYP3A4, respectively. The heatmaps show both the fraction of genes in a given AD gene set (columns, *x* axis) that interact with a given mediator. The number of interacting AD genes is also indicated as text inside the heatmap’s cells. (B) All three edge-long paths between the three drugs and knowledge-based AD genes. For mediator genes thick, color edges mark paths formed by target → mediator → AD gene. For paths involving RGS4, the color codes for mediator genes are as follows. Orange: EGFR (Epidermal growth factor receptor), purple: CALM1 (Calmodulin 1), gray: COPB1 (Coatomer subunit beta), also gray: GNAO1 (Guanine nucleotide-binding protein G(o) subunit alpha).

RGS4 interacts with AD genes either directly (Table S3) or indirectly through mediator genes (Fig. 4). These interactions implicate RGS4-dependent mechanisms involving two ErbB signaling pathways: MAPK/ERK via EGFR (ErbB1) and PI3K/Akt/mTOR via ERBB3. EGFR and ERBB3 regulate proliferation, differentiation, and maturation in the nervous system and other tissues and are central in cancer biology [33, 34]. Although less is known for ERBB3 in neurodegenerative disease [35], broad evidence supports EGFR dysregulation in AD, Parkinson’s disease, and ALS [36, 37]. Our cell-based findings that arundine and TUDCA lower neuroin-flammation, enhance A*β* clearance, and reduce A*β* release are consistent with these pathways.

Several EGFR inhibitors, including blood-brain-penetrant agents, have FDA approval in oncology. This converges with our finding that drugs approved for glioblastoma, carcinomas, and other neoplasms are rediscovered by an AD-focused network screen, supporting repurposing potential of anti-cancer EGFR-pathway drugs in AD [37].

Our Bayesian regression framework helped us test protective hypotheses directly at biologically plausible effect sizes rather than relying only on null-hypothesis p-values. The finding that TUDCA is promising in AD is also consistent with prior results linking bile-acid metabolism to amyloid burden, brain atrophy, and vascular dementia risk [38].

Computational drug repurposing is a rapidly evolving field with diverse approaches and increasingly rich multi-omic inputs [3], including recent AD applications [4, 5]. A key strength of our screen is use of multiple complementary AD risk gene sets, including recent fine-mapping and iPSC-derived single-cell datasets. The systems-level network-pharmacology framework integrates drug-target and gene-gene networks to identify plausible drugs acting on putative causal disease pathways.

Important limitations include incompleteness and noise in underlying drug-target and gene-gene interaction networks, and our restriction to FDA-approved or latestage clinical compounds. Future work should expand to newer chemical entities and updated interaction networks.

In summary, we applied a network pharmacology strategy that prioritizes RGS4-linked drug mechanisms and generated experimental evidence that TUDCA and arundine are promising repurposing candidates for AD.

## Acknowledgments

We thank Yang An for discussions on statistical methodology and the NIH High Performance Computing group for providing us with the Biowulf cluster.

## Author Contributions

M.T conceived the study overall and secured funding. A.J and M.T conceived the computational part of the study. A.J designed, implemented and carried out the computational part of the study. E.W and H.K.I supported the computational part of the study using S-PrediXcan. M.T, V.R.V and T.L designed and managed the experimental part of the study. T.L performed experiments (cell-based assays). A.J developed Bayesian statistical tools for, and analyzed, experimental data. A.J prepared the first draft of the manuscript. A.J, M.T and V.R.V reviewed and edited the manuscript. All authors have read and approved the final draft and take full responsibility of its content, including the accuracy of the data and the fidelity of the trial to the registered protocol and its statistical analysis.

## 5 Tables

**Table 1.** AD risk gene sets used as inputs to drug repurposing screens of this study. For the TWAS2+ gene set we combined gene sets from the prior published TWAS with the gene set from our own TWAS (Methods). The genes in each set are listed in Table S1.

| AD risk gene set | Size | Description | Reference |
| --- | --- | --- | --- |
| knowledge | 27 | curated AD risk genes from the DISEASES database | [39] |
| TWAS2+ | 32 | genes supported by $\geq 2$ AD-TWAS/PWAS | [9–13, 15] |
| agora2+ | 64 | genes supported by $\geq 2$ Agora studies | https://agora.adknowledgeportal.org |
| AD DE APOE3-APOE3 | 277 | AD vs control DEGs: APOE3/APOE3 background | [5] |
| AD DE APOE4-APOE4 | 274 | AD vs control DEGs: APOE4/APOE4 background | [5] |
| APOE3-4 DE neuron | 46 | APOE4 vs APOE3 DEGs: iPSC-derived neurons | [16] |
| APOE3-4 DE astrocyte | 128 | APOE4 vs APOE3 DEGs: iPSC-derived astrocytes | [16] |
| APOE3-4 DE microglia | 140 | APOE4 vs APOE3 DEGs: iPSC-derived microglia | [16] |

**Table 2.**
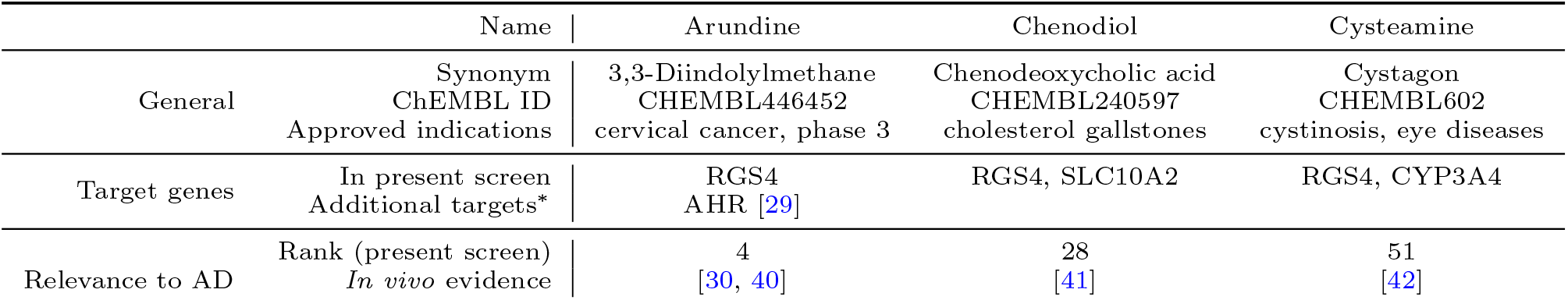
Selected drugs ranked high by the present computational screening of 2413 drugs from ChEMBL. ChEMBL Ranking was established based on a given drugs permeability across the blood-brain barrier, network proximity of drug-target and AD risk genes and their prior evidence for roles in AD pathogenesis. Notes: Targets in the present screen were taken from ChEMBL filtering for human experiments. ^∗^“Additional targets” refer to those we found manually in the biomedical literature and are not in the drug-target network used for our screen.

