## Supplemental Information for "Network pharmacology-based discovery and experimental validation of novel drug repurposing candidates in Alzheimer’s Disease"

#### Supplemental Methods

##### Previous and novel TWAS on AD

We obtained AD risk gene sets from multiple TWAS. Besides collecting AD risk genes from previous transcriptome- and proteome-wide association studies (TWAS, PWAS) we conducted our own applying the S-MultiXcan approach [1] to the two largest AD-GWAS to date [2, 3]. S-MultiXcan borrows statistical strength from multiple tissue types; we used a predictive model of gene expression built on all tissues of the GTEx data set or on only brain tissues. We conservatively took the top 30 genes and investigated their agreement with top gene sets from seven previous TWAS and one previous PWAS (see Data file 1) listing genes contained in each set). Hierarchical clustering of studies based on shared genes (Fig. S2) showed a fairly low agreement for all pairs of studies. The most similar study pair was our present TWAS and the fine mapping of [18] despite several methodological differences between these two studies, which helps validate our novel TWAS results. As might be expected, the least similarity was found between the only PWAS in our collection and all the other studies.

##### A$\beta$ clearance

For A$\beta$ clearance assay, 20,000 BV2 cells per well (uncoated 96-well plates) were plated out. After changing cells to treatment medium, drug compounds were administered 1 hour before A$\beta$ stimulation (Bachem, 4061966; final concentration in well, 200 ng/ml; dilutions in medium). Cells treated with vehicle and cells treated with A$\beta$ alone served as controls. After 3 hours of A$\beta$ stimulation, cell supernatants were collected for the A$\beta$ measurement and cells were carefully washed twice with PBS and thereafter lysed in 35 $\mu$l of cell lysis buffer (50 mM tris-HCl, pH 7.4, 150 mM NaCl, 5 mM EDTA, and 1% SDS) supplemented with protease inhibitors. Supernatants and cell lysates were analyzed for human A$\beta$42 with the MSD V-PLEX Human A$\beta$42 Peptide (6E10) Kit (K151LBE, Mesoscale Discovery). The immune assay was carried out according to the manual, and plates were read on MESO QuickPlex SQ 120. For a separate set of experiments iPSC derived microglia from an adult human AD patient were used (FUJIFILM Cellular Dynamics Inc, CatNo 1212, Female, APOE 4/4, Alzheimer's Disease) and maintained according to the provided manual. Cells were seeded at a density of 10,000 cells per well in 384 well plate in the kit provided media. Drug treatment was applied 1 hour before A$\beta$1-42 stimulation (Bachem 4061966; final concentration in well: 4 $\mu$M (dilutions in medium) labelled with pHrodo Red (Thermo Fisher, P36011). Cells treated with vehicle (H2O) and cells treated with A$\beta$1-42 alone served as controls. After 4 h and 24 h of A$\beta$1-42 stimulation, cell supernatants were collected for the A1-42 measurement. Supernatants were analyzed for human A$\beta$1-42 with MSD V-PLEX Human A$\beta$42 Peptide (6E10) Kit (K151LBE, Mesoscale Discovery). The immune assay was carried out according to the manufacturers manual and plates were read on the MESO QuickPlex SQ 120.

#### A$\beta$ clearance

H4-hAPP cells were cultivated in Opti-MEM supplemented with 10% FCS, 1% penicillin/streptomycin, hygromycin B (200 $\mu$g/ml), and blasticidin S (2.5 $\mu$g/ml). H4-hAPP cells were seeded into 96-well plates (2 × 104 cells per well). On the next day, cells in 96-well plates were treated with compounds, reference item [400 nM N-[N-(3,5-difluorophenacetyl-l-alanyl)]-S-phenylglycine t-butyl ester (DAPT)], or vehicle. Twenty-four hours later, supernatants were collected for further A$\beta$ measurements by MSD [V-PLEX A$\beta$ Peptide Panel 1 (6E10) Kit, K15200E, Mesoscale Discovery].

#### Tau phosphorylation

SH-SY5Y-hTau441(V337M/R406W) cells were maintained in culture medium [DMEM medium, 10% FCS, 1% nonessential amino acids (NEAA), 1% l-glutamine, gentamycin (100 $\mu$g/ml), and geneticin G-418 (300 $\mu$g/ml)] and differentiated with 10 $\mu$M retinoic acid for 5 days changing medium every 2 to 3 days. Before the treatment, cells were seeded onto 24-well plates at a cell density of 2 × 105 cells per well on day one of in vitro culture (DIV1). Drug compounds were applied on DIV2. After 24 hours of incubation (DIV3), cells on 24-well plates were harvested in 60 $\mu$l of RIPA buffer [50 mM tris (pH 7.4), 1% NP-40, 0.25% Na-deoxycholate, 150 mM NaCl, 1 mM EDTA supplemented with freshly added 1 $\mu$M NaF, 0.2 mM Na-orthovanadate, 80 $\mu$M glycerophosphate, protease (Calbiochem), and phosphatase (Sigma-Aldrich) inhibitor cocktail]. Protein concentration was determined by BCA assay (Pierce, Thermo Fisher Scientific), and samples were adjusted to a uniform total protein concentration. Total tau and phosphorylated tau were determined by immunosorbent assay from Mesoscale Discovery [Phospho(Thr231)/Total Tau Kit K15121D, Mesoscale Discovery].

#### LPS-induced neuroinflammation

The murine microglial cell line BV2 was cultivated in Dulbecco's modified Eagle's medium (DMEM) medium supplemented with 10% fetal calf serum (FCS), 1% penicillin/streptomycin, and 2 mM l-glutamine (culture medium). For LPS stimulation assay, 5000 BV2 cells per well (uncoated 96-well plates) were plated out and the medium was changed to treatment medium (DMEM, 5% FCS, and 2 mM l-glutamine). After changing cells to treatment medium, drug compounds were administered 1 hour before LPS stimulation [Sigma-Aldrich; L6529; 1 mg/ml stock in ddH2O; final concentration in well, 100 ng/ml (dilutions in medium)]. Cells treated with vehicle, cells treated with LPS alone, and cells treated with LPS plus reference item (dexamethasone, 10 $\mu$M; Sigma-Aldrich, D4902) served as controls. After 24 hours of stimulation, cell supernatants were collected for the cytokine measurement (V-PLEX Proinflammatory Panel 1 Mouse Kit, K15048D, Mesoscale) and cells were subjected to 3-(4,5-dimethylthiazol-2-yl)-2,5-diphenyltetrazolium bromide (MTT) assay.

#### Neurite outgrowth and neurogenesis

Primary hippocampal neurons were prepared from E18.5 timed pregnant C57BL/6JRccHsd mice as previously described. Cells were seeded in poly-D-lysine pre-coated 96-well plates at a density of 2.6 × 104 cells/well in medium (Neurobasal, 2% B-27, 0.5 mM glutamine, 25 M glutamate, 1% Penicillin-Streptomycin). Directly on DIV1, the drug of interest or VC was applied. On DIV2, 10 $\mu$M Bromodeuxyuridine (BrdU; B5002 Sigma Aldrich) was added and cells were fixed after additional 24h. Cells were permeabilized with 0.1% Triton-X and incubated with primary Beta Tubulin Isotype III (T8660, Sigma Aldrich) and BrdU antibodies (MAS25°C, AHrlan-Sera Lab) overnight at 4°C. Afterwards, cells were washed two times with PBS and incubated with fluorescently labelled secondary antibodies and DAPI for 1.5 h at room temperature in the dark. Cells were rinsed three times with PBS and imaged with the Cytation 5 Multimode reader (BioTek) at 10 × magnification (six images per well). BrdU-positive cells were counted as a marker of neurogenesis and Beta Tubulin Isotype III signal was used for macro-based quantification of neurite outgrowth.

#### Trophic factor withdrawal

Primary cortical neurons from embryonic day 18 (E18) C57Bl/6 mice were prepared as previously described. On the day of preparation (DIV1), cortical neurons were seeded on poly-d-lysine precoated 96-well plates at a density of 3 × 104 cells per well. Every 4 to 6 days, a half medium exchange using full medium (Neurobasal, 2% B-27, 0.5 mM glutamine, and 1% penicillin-streptomycin) was carried out. On DIV8, a full medium exchange to B-27 free medium (Neurobasal, 0.5 mM glutamine, and 1% penicillin-streptomycin) was performed and drug compounds were applied thereafter. The experiment was carried out with six technical replicates per condition, and vehicle-treated cells served as control. After 28 hours on B-27-free medium, cells were subjected to YO-PRO/propidium iodide (PI) and MTT as well as lactate dehydrogenase (LDH) assay.

### Supplemental Discussion

We and others have shown that bile acid synthesis, including that of the primary bile acid, chenodiol, or Chenodeoxycholic acid (CDCA), is linked to AD pathology and dementia risk [4–8]. Moreover, chenodiol has been shown to be neuroprotective in Huntingtons disease [9]. One of the chenodiol targets that our analysis implicated is the RGS4 gene (regulator of G-protein signaling 4). RGS4 is notable because it is also targeted by arundine, another of our selected top-ranking drugs. arundine, also known as 3,3-diindolylmethane, is the dimeric product of the natural product indole-3-carbinol. While arundine has been mostly investigated in the context of drug-resistant tumors [10], its close analogs have been found to cross the blood-brain barrier and protect mice from 1-methyl-4-phenyl-1,2,3,6-tetrahydropyridine (MPTP)-induced neurotoxicity and neurodegeneration [11]. Moreover, arundine was shown to inhibit oxidative stress-induced apoptosis in hippocampal neuronal cells [12] and protected primary hippocampal cell cultures from ischemia-induced apoptosis and autophagy [13]. The latter effect was found to depend on arundine binding to the Aryl Hydrocarbon Receptor, product of the AHR gene, [13].

Our third selected drug was cysteamine. Cysteamine, whose targets also include RGS4, is a derivative of the amino acid cysteine. Cysteamine can traverse the blood brain barrier and is an approved therapy for cystinosis (MeSH:D005128). More recently it has been studied as a potential drug in Huntingtons and Parkinsons disease due to its neuroprotective activity [14, 15].

### Supplemental Tables

| | | | |
|---|---|---|---|
| knowledge | 27 | curated AD risk genes from the DISEASES database | [16] |
| TWAS2+ | 32 | genes supported by $\geq 2$ AD-TWAS/PWAS | [2, 17–20] |
| agora2+ | 64 | genes supported by $\geq 2$ Agora studies | * |
| AD DE APOE3-APOE3 | 277 | AD vs control DEGs: APOE3/APOE3 background | [21] |
| AD DE APOE4-APOE4 | 274 | AD vs control DEGs: APOE4/APOE4 background | [21] |
| APOE3-4 DE neuron | 46 | APOE4 vs APOE3 DEGs: iPSC-derived neurons | [22] |
| APOE3-4 DE astrocyte | 128 | APOE4 vs APOE3 DEGs: iPSC-derived astrocytes | [22] |
| APOE3-4 DE microglia | 140 | APOE4 vs APOE3 DEGs: iPSC-derived microglia | [22] |

**Table S 1** AD risk gene sets used as inputs to drug repurposing screens. Abbreviations: DEdifferentially expressed; DEGsdifferentially expressed genes; TWAS (PWAS)transcriptome(proteome)-wide association study; iPSCinduced pluripotent stem cell. *https://agora.adknowledgeportal.org

| Name | arundine | chenodiol | cysteamine |
|---|---|---|---|
| Synonym | 3,3-Diindolylmethane | chenodeoxycholic acid | cystagon |
| ChEMBL ID | CHEMBL446452 | CHEMBL240597 | CHEMBL602 |
| Approved indications | cervical cancer, phase | cholesterol gallstones | cystinosis, eye diseases |
| Targets (in drug-target network) | RGS4 | RGS4, SLC10A2 | RGS4, CYP3A4 |
| Targets (additional) | AHR [13] | | |
| Rank (present screen) | 4 | 28 | 51 |
| Prior in vivo evidence | [11] | [9] | [23] |

**Table S 2** Selected drugs ranked high by the present computational screening of 2413 drugs from ChEMBL. ChEMBL. Ranking was established based on a given drugs permeability across the blood-brain barrier, network proximity of drug-target and AD risk genes and their prior evidence for roles in AD pathogenesis. Notes: Targets in the present screen were taken from ChEMBL filtering for human experiments. ∗"Additional targets" refer to those we identified in the biomedical literature and were not in the drug-target network used for our screen. In vivo evidence refers to studies suggesting a neuroprotective role in preclinical animal models of AD.

| | | target | | |
|---|---|---|---|---|
| | | CYP3A4 | RGS4 | SLC10A2 |
| shortest path length | AD risk gene set | AD risk genes | | |
| 1 | AD DE APOE3-APOE3 | - | RGS4 | - |
| 1 | AD DE APOE4-APOE4 | - | RGS4 | - |
| 2 | agora2+ | - | ERBB3 | - |
| 2 | AD DE APOE3-APOE3 | - | GNAI2 | - |
| 2 | AD DE APOE4-APOE4 | - | CALM1; GNAI2 | - |
| 2 | APOE3-4 DE neuron | - | PLCB1 | - |

**Table S 3** Targets of the drugs Arundine, Cysteamine or Chenodiol: CYP3A4, RGS4 and SLC10A2 and network paths going from the drugs through the targets to AD genes in some AD gene set. When the target is itself an AD gene, the shortest path length is 1. When the target directly interacts with an AD gene, the path length is 2. We see that RGS4 is the only target that fulfils either criterion.

| experiment | assay | desired effect |
|---|---|---|
| LPS neuroinflammation (BV2 cells) | IFN-$\gamma$ | decrease |
| | IL-10 | decrease |
| | IL-12p70 | decrease |
| | IL-1$\beta$ | decrease |
| | IL-2 | decrease |
| | IL-4 | decrease |
| | IL-5 | decrease |
| | IL-6 | decrease |
| | KC/GRO | decrease |
| | TNF-$\alpha$ | decrease |
| A$\beta$ toxicity (primary neurons) | MTT | increase |
| A$\beta$ release (H4 cells) | A$\beta$38 | decrease |
| | A$\beta$40 | decrease |
| | A$\beta$42 | decrease |
| A$\beta$ clearance (BV2 cells) | A$\beta$42 SN | decrease |
| | A$\beta$42 Ly | increase |
| Trophic factor withdrawal (primary neurons) | PI | decrease |
| | YOPRO | decrease |
| | MTT | increase |
| | LDH | decrease |
| Tau phosphorylation | Tau | increase |
| | pTau (T231) | decrease |
| | pT/T ratio | decrease |
| Neurite outgrowth (primary neurons) | $\sum$ neurite area | increase |
| | branch points | increase |
| | neurogenesis | increase |
| | longest neurite | increase |
| A$\beta$ clearance (iPSC) | pHrodo-4h | increase |
| | supernatant | decrease |
| LPS neuroinflammation (iPSC) | IL-1$\beta$ | decrease |
| | IL-6 | decrease |
| | IL-8 | decrease |
| | MTT | decrease |
| | TNF-$\alpha$ | decrease |

**Table S 4** Desired, neuroprotective, effect for each assay. Desired effect is the direction of effect of an ideal, hypothetical, neuroprotective drug on a given assay.

### Supplemental Figures

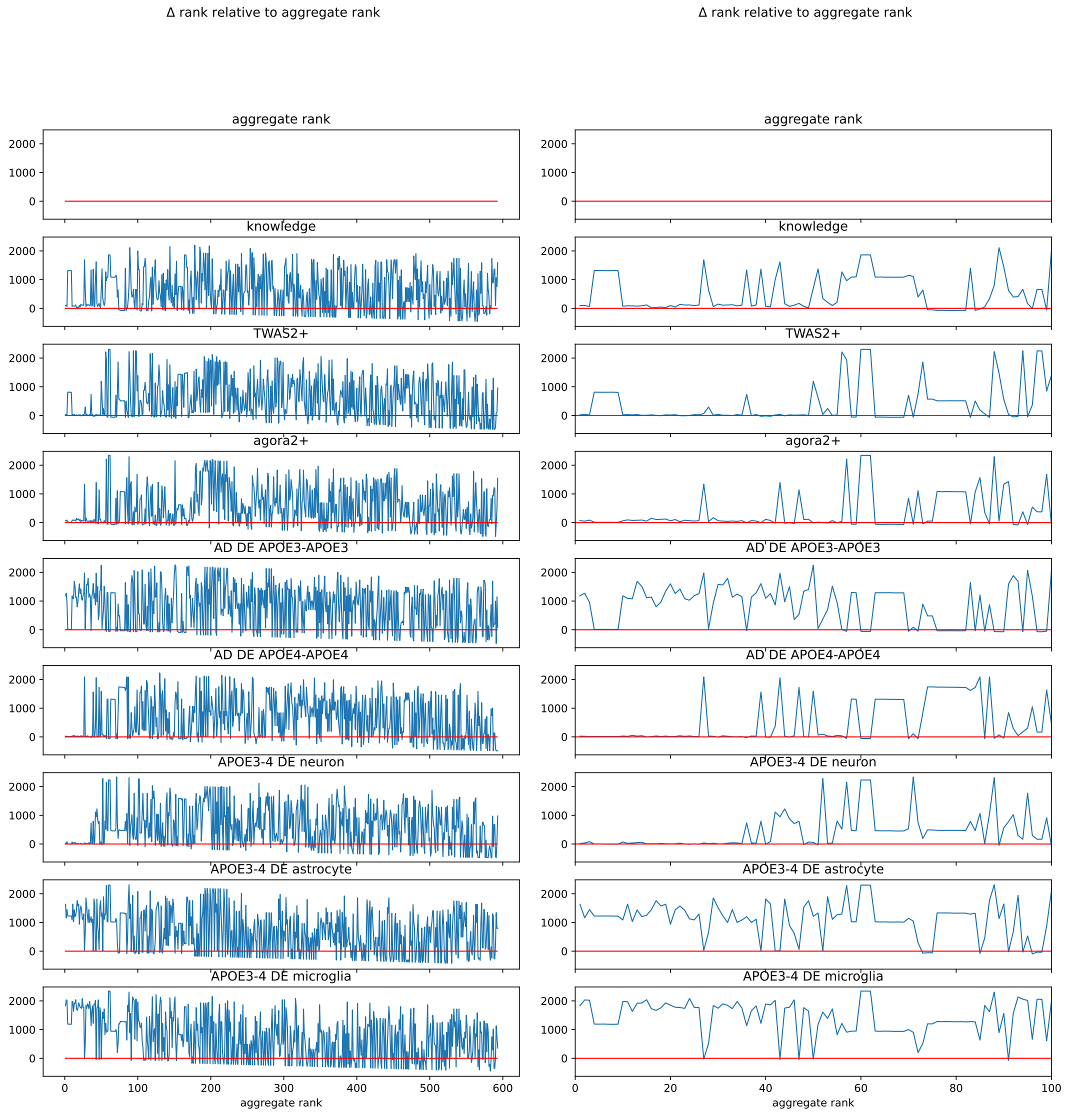


**Figure S 1** Difference of drug rank between the final, aggregated list and the list resulting from each network proximity based drug screen.

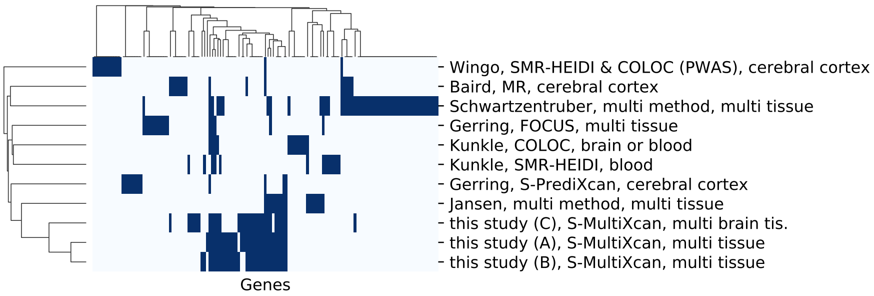


**Figure S 2** Similarity of TWA/PWA studies on AD in terms of shared genes.

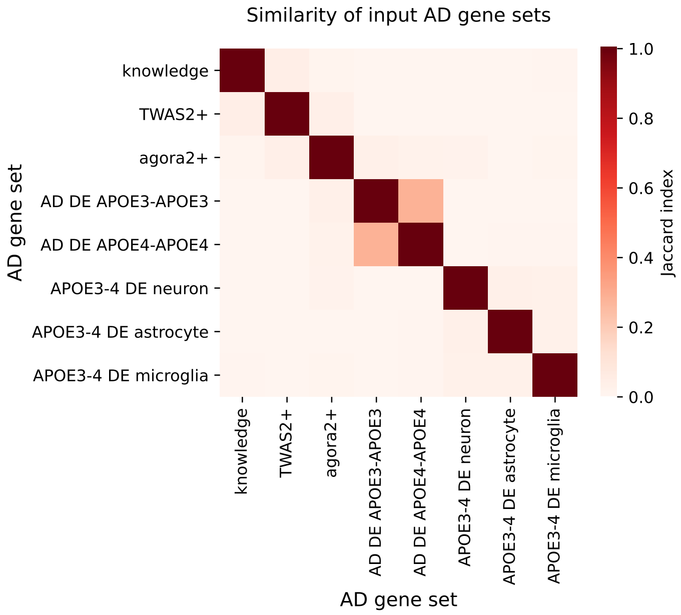


**Figure S 3** Eight AD risk gene sets were input to the workflow. Jaccard indices quantifying shared genes show that the sets are highly dissimilar to each other.

### Supplemental Data Files

Data file 1. Genes of the AD risk gene sets used as inputs to the present computational drug screen.

Data file 2. The 2413 drugs ranked according to their network proximity to each of the eight AD risk gene sets used as input. The drugs final, aggregate rank is also shown as well as their ChEMBL ID, standard InChI, indication class, and blood-brain-barrier permeability taken (if available) from the BBB database (Meng et al. 2021). Moreover, the UniProt name of each drugs targets is also indicated.

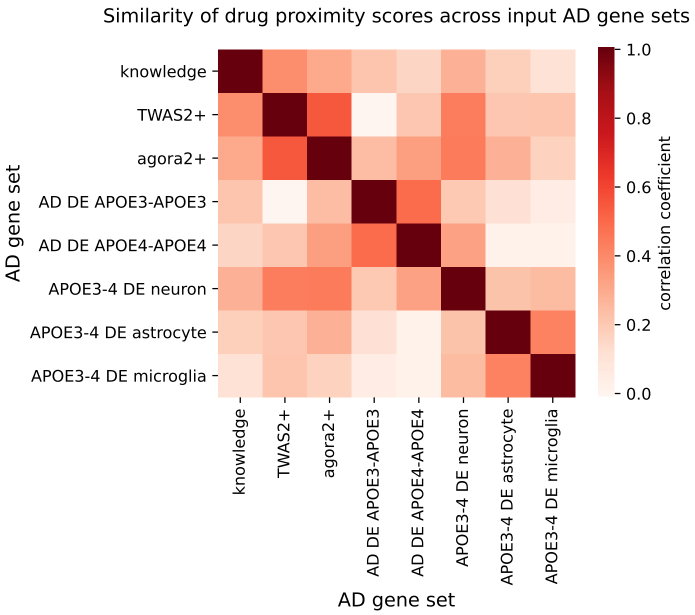


**Figure S 4** The network proximity scores of all screened drugs are fairly similar to each other across the different input AD risk gene sets, in spite of the marked dissimilarity among those.

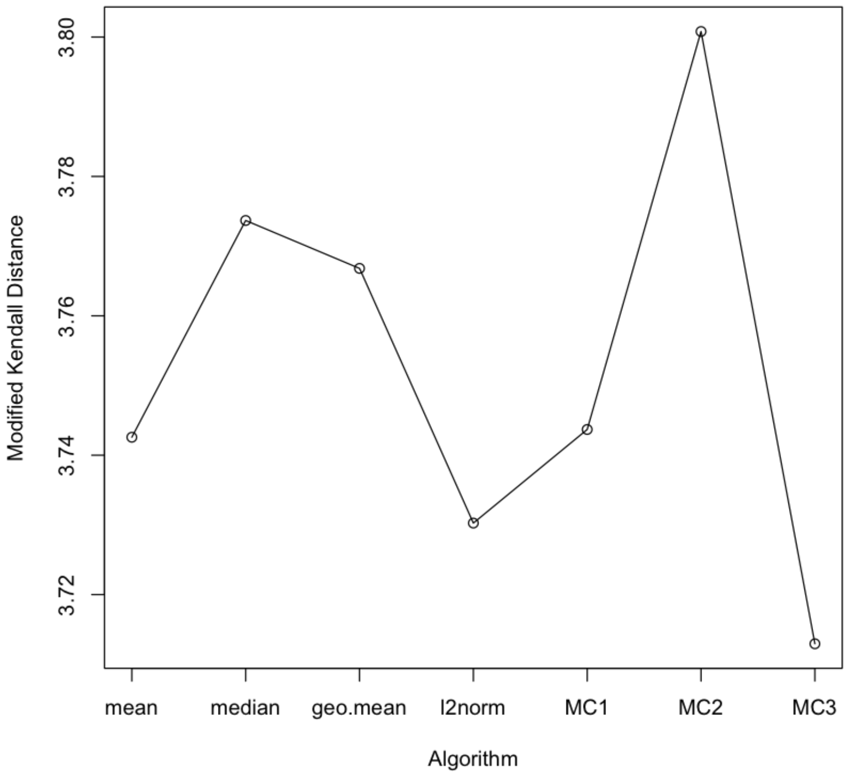


**Figure S 5** Several rank aggregation algorithms' performance assessed by modified Kendall distance.

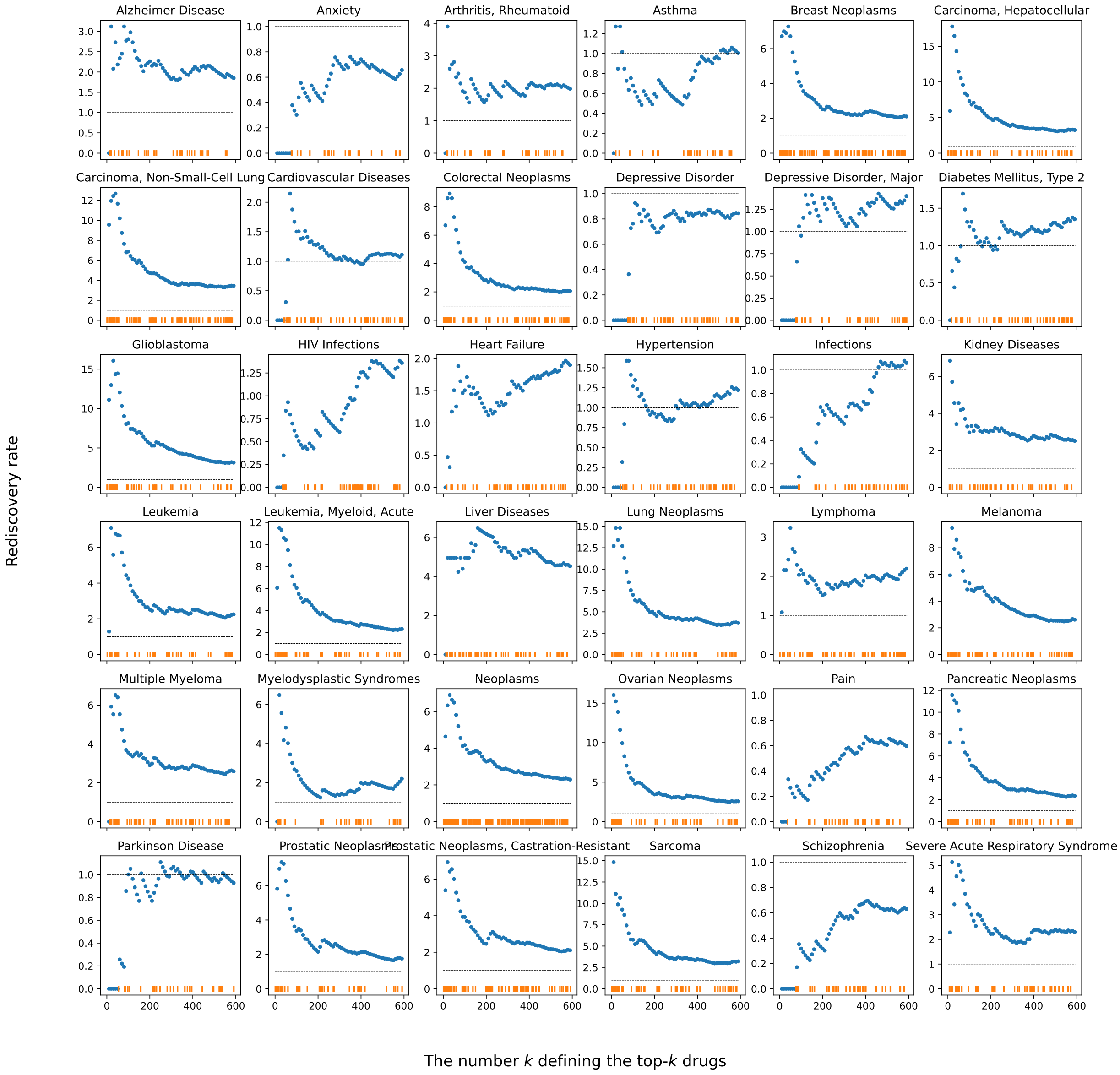


**Figure S 6** Network proximity-based rediscovery of drugs in phase 1–4 clinical trials for AD and 35 other disease indications. Disease indications written on top of individual plots. Orange ticks: drugs that are in phase 1 or more advanced clinical study for the given indication. Blue dots: rediscovery rate (Eq. 3, Methods
) for the top-$k$ drugs, ranked by network proximity, relative to that for the bottom-1808 drugs. Rediscovery rate of $> 1$ means that top-ranked drugs tend to be those that are in phase 1–4 clinical trials for the given indication; these drugs are marked by orange symbols above the $x$ axes.

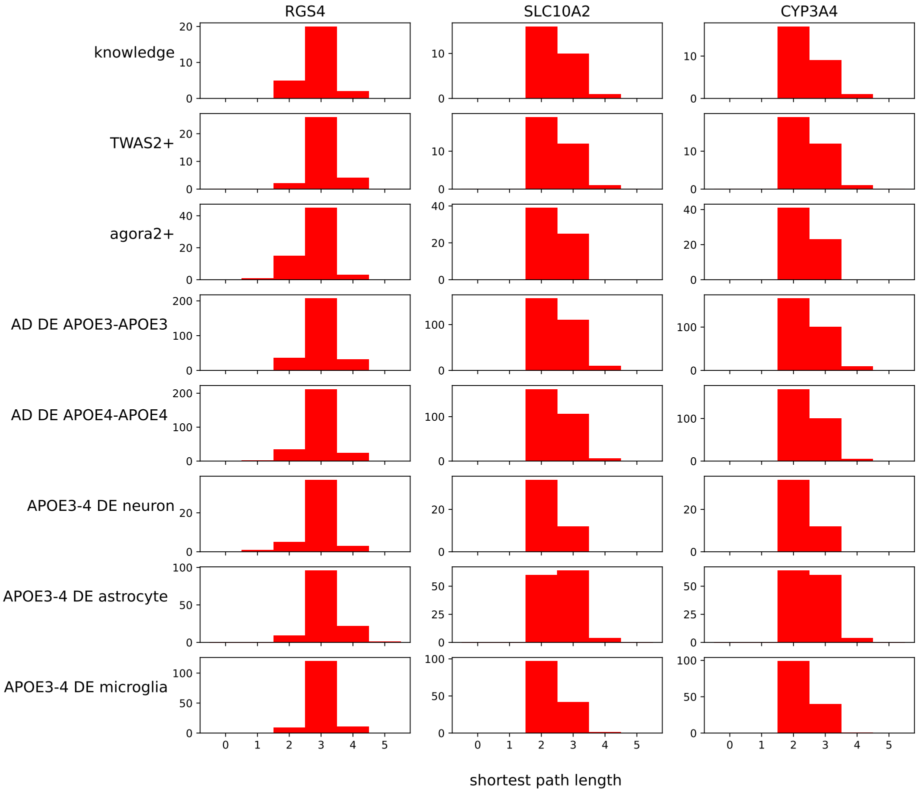


**Figure S 7** Shortest path length distribution from a given drug target (columns) to a given AD risk gene set (rows) across all genes in the set.

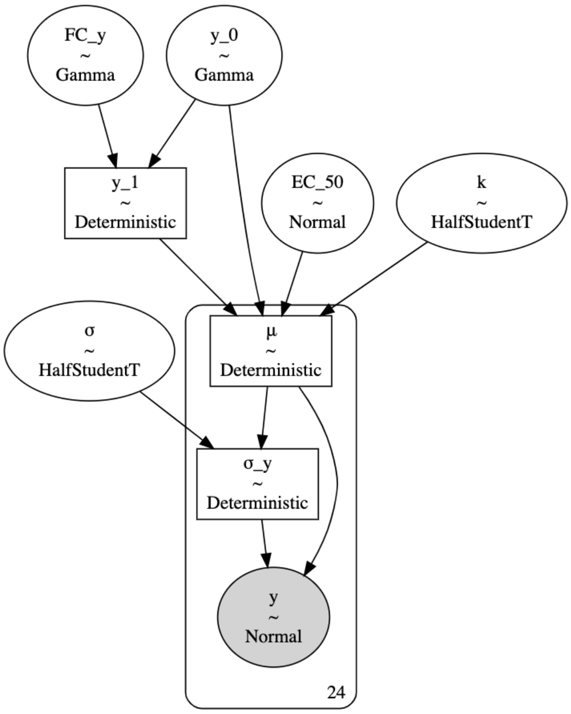


**Figure S 8** Dependency graph of the Bayesian nonlinear regression model used to fit dose-response ($x$–$y$) data from cell-based assays. For more details see Eq.4 in Methods. This model is named "sigmoid 2" in https://github.com/attilagk/CTNS-notebook/blob/main/src/cellbayesassay.py (see the definition of the sample_sigmoid_2 function therein).

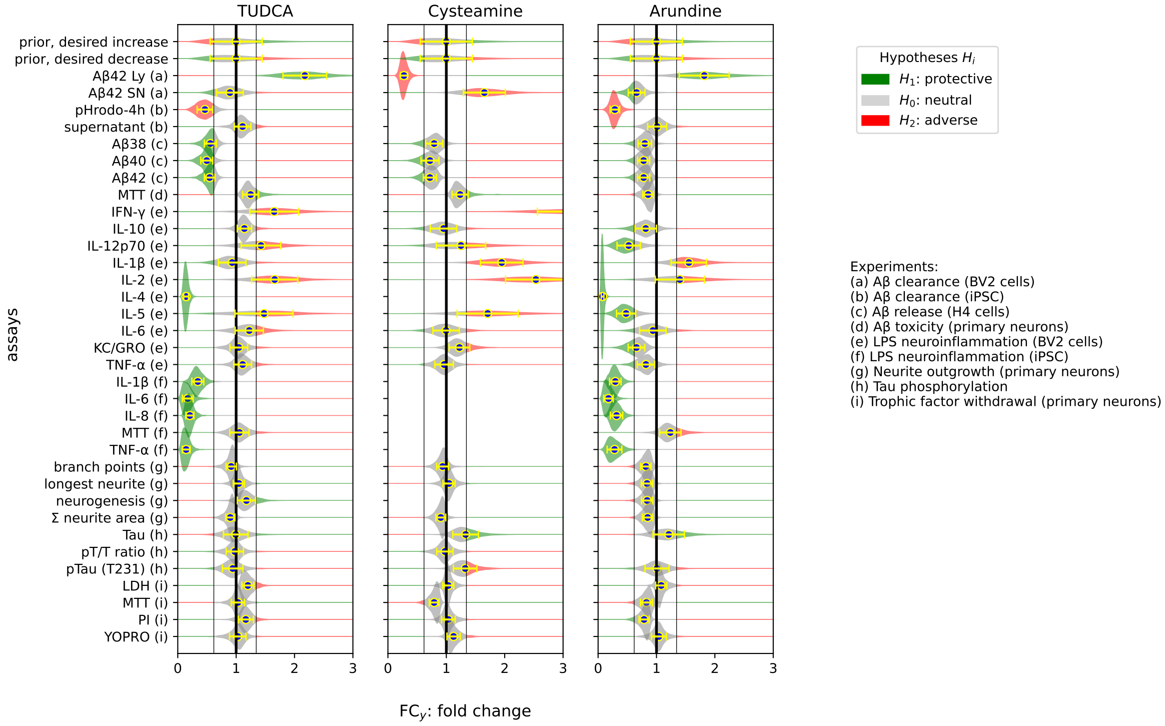


**Figure S 9** Posterior probability density of drug-induced fold change in bioactivity across three candidate drugs and 33 cell-based assays. Blank (white) slots indicate either that the experiment was not performed on the drug or that model fit was poor. One or more assays constitute an experiment (see parenthesized one-letter codes and the right center legend). Each posterior density has a green, gray and red component corresponding to protective, neutral and adverse drug effect given the direction of the desired drug effect and fold-change thresholds $t_1, t_2$ (see Fig. S**??**). The posterior mean (and standard deviation) of fold change is shown as a yellow circle (and yellow error bars).

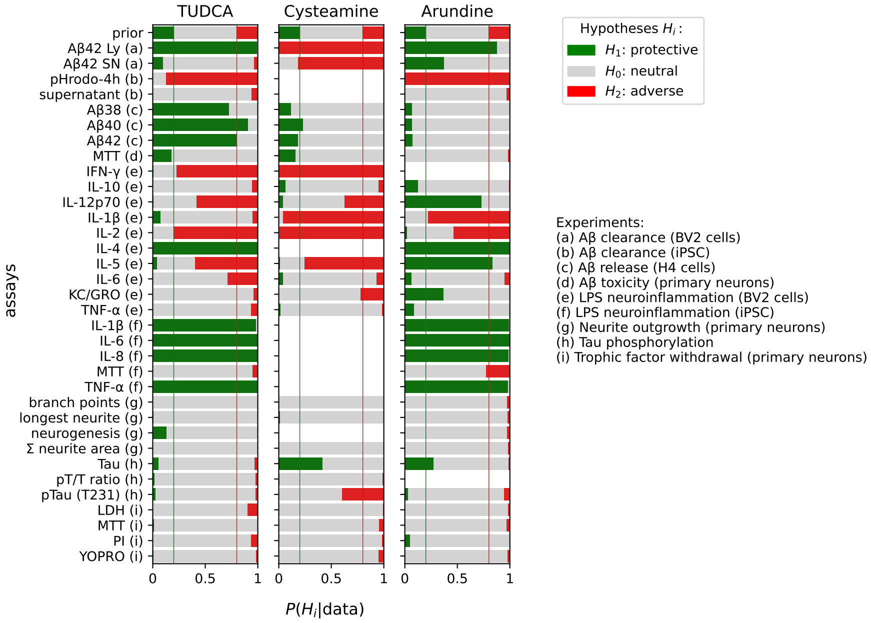


**Figure S 10** Posterior probability of protective, neutral and adverse effect for Arundine, Cysteamine and TUDCA.

#### References


[1] Barbeira, A.N., Dickinson, S.P., Bonazzola, R., Zheng, J., Wheeler, H.E., Torres, J.M., Torstenson, E.S., Shah, K.P., Garcia, T., Edwards, T.L., Stahl, E.A., Huckins, L.M., Aguet, F., Ardlie, K.G., Cummings, B.B., Gelfand, E.T., Getz, G., Hadley, K., Handsaker, R.E., Huang, K.H., Kashin, S., Karczewski, K.J., Lek, M., Li, X., MacArthur, D.G., Nedzel, J.L., Nguyen, D.T., Noble, M.S., Segr, A.V., Trowbridge, C.A., Tukiainen, T., Abell, N.S., Balliu, B., Barshir, R., Basha, O., Battle, A., Bogu, G.K., Brown, A., Brown, C.D., Castel, S.E., Chen, L.S., Chiang, C., Conrad, D.F., Damani, F.N., Davis, J.R., Delaneau, O., Dermitzakis, E.T., Engelhardt, B.E., Eskin, E., Ferreira, P.G., Frsard, L., Gamazon, E.R., Garrido-Martn, D., Gewirtz, A.D.H., Gliner, G., Gloudemans, M.J., Guigo, R., Hall, I.M., Han, B., He, Y., Hormozdiari, F., Howald, C., Jo, B., Kang, E.Y., Kim, Y., Kim-Hellmuth, S., Lappalainen, T., Li, G., Li, X., Liu, B., Mangul, S., McCarthy, M.I., McDowell, I.C., Mohammadi, P., Monlong, J., Montgomery, S.B., Muoz-Aguirre, M., Ndungu, A.W., Nobel, A.B., Oliva, M., Ongen, H., Palowitch, J.J., Panousis, N., Papasaikas, P., Park, Y., Parsana, P., Payne, A.J., Peterson, C.B., Quan, J., Reverter, F., Sabatti, C., Saha, A., Sammeth, M., Scott, A.J., Shabalin, A.A., Sodaei, R., Stephens, M., Stranger, B.E., Strober, B.J., Sul, J.H., Tsang, E.K., Urbut, S., Bunt, M., Wang, G., Wen, X., Wright, F.A., Xi, H.S., Yeger-Lotem, E., Zappala, Z., Zaugg, J.B., Zhou, Y.-H., Akey, J.M., Bates, D., Chan, J., Chen, L.S., Claussnitzer, M., Demanelis, K., Diegel, M., Doherty, J.A., Feinberg, A.P., Fernando, M.S., Halow, J., Hansen, K.D., Haugen, E., Hickey, P.F., Hou, L., Jasmine, F., Jian, R., Jiang, L., Johnson, A., Kaul, R., Kellis, M., Kibriya, M.G., Lee, K., Li, J.B., Li, Q., Li, X., Lin, J., Lin, S., Linder, S., Linke, C., Liu, Y., Maurano, M.T., Molinie, B., Montgomery, S.B., Nelson, J., Neri, F.J., Oliva, M., Park, Y., Pierce, B.L., Rinaldi, N.J., Rizzardi, L.F., Sandstrom, R., Skol, A., Smith, K.S., Snyder, M.P., Stamatoyannopoulos, J., Stranger, B.E., Tang, H., Tsang, E.K., Wang, L., Wang, M., Van Wittenberghe, N., Wu, F., Zhang, R., Nierras, C.R., Branton, P.A., Carithers, L.J., Guan, P., Moore, H.M., Rao, A., Vaught, J.B., Gould, S.E., Lockart, N.C., Martin, C., Struewing, J.P., Volpi, S., Addington, A.M., Koester, S.E., Little, A.R., Brigham, L.E., Hasz, R., Hunter, M., Johns, C., Johnson, M., Kopen, G., Leinweber, W.F., Lonsdale, J.T., McDonald, A., Mestichelli, B., Myer, K., Roe, B., Salvatore, M., Shad, S., Thomas, J.A., Walters, G., Washington, M., Wheeler, J., Bridge, J., Foster, B.A., Gillard, B.M., Karasik, E., Kumar, R., Miklos, M., Moser, M.T., Jewell, S.D., Montroy, R.G., Rohrer, D.C., Valley, D.R., Davis, D.A., Mash, D.C., Undale, A.H., Smith, A.M., Tabor, D.E., Roche, N.V., McLean, J.A., Vatanian, N., Robinson, K.L., Sobin, L., Barcus, M.E., Valentino, K.M., Qi, L., Hunter, S., Hariharan, P., Singh, S., Um, K.S., Matose, T., Tomaszewski, M.M., Barker, L.K., Mosavel, M., Siminoff, L.A., Traino, H.M., Flicek, P., Juettemann, T., Ruffier, M., Sheppard, D., Taylor, K., Trevanion, S.J., Zerbino, D.R., Craft, B., Goldman, M., Haeussler, M., Kent, W.J., Lee, C.M., Paten, B., Rosenbloom, K.R., Vivian, J., Zhu, J., Nicolae, D.L., Cox, N.J., Im, H.K., Consortium, G., Laboratory, D.A..C.C.L.-A.W.G., Group, S.M.g.-A.W., groups, E.G., Fund, N.I.H.C., NIH/NCI, NIH/NHGrI, NIH/NIMH, NIH/NIDA, Site-NDrI, B.C.S.,

Site-rPCI, B.C.S., resource-VArI, B.C., Miami Brain Endowment Bank, B.B.r.-U., Management, L.B.-P., Study, E.L.S.I., Visualization-EBI, G.B.D.I.., Genome Browser Data Integration & Visualization-UCSC Genomics Institute, U.o.C.S.C.: Exploring the phenotypic consequences of tissue specific gene expression variation inferred from gwas summary statistics. Nature Communications **9**(1), 1825 (2018) https://doi.org/10.1038/s41467-018-03621-1

[2] Schwartzentruber, J., Cooper, S., Liu, J.Z., Barrio-Hernandez, I., Bello, E., Kumasaka, N., Young, A.M.H., Franklin, R.J.M., Johnson, T., Estrada, K., Gaffney, D.J., Beltrao, P., Bassett, A.: Genome-wide meta-analysis, fine-mapping and integrative prioritization implicate new alzheimer's disease risk genes. Nature genetics **53**(33589840), 392–402 (2021) https://doi.org/10.1038/s41588-020-00776-w

[3] Wightman, D.P., Jansen, I.E., Savage, J.E., Shadrin, A.A., Bahrami, S., Holland, D., Rongve, A., Brte, S., Winsvold, B.S., Drange, O.K., Martinsen, A.E., Skogholt, A.H., Willer, C., Brthen, G., Bosnes, I., Nielsen, J.B., Fritsche, L.G., Thomas, L.F., Pedersen, L.M., Gabrielsen, M.E., Johnsen, M.B., Meisingset, T.W., Zhou, W., Proitsi, P., Hodges, A., Dobson, R., Velayudhan, L., Sealock, J.M., Davis, L.K., Pedersen, N.L., Reynolds, C.A., Karlsson, I.K., Magnusson, S., Stefansson, H., Thordardottir, S., Jonsson, P.V., Snaedal, J., Zettergren, A., Skoog, I., Kern, S., Waern, M., Zetterberg, H., Blennow, K., Stordal, E., Hveem, K., Zwart, J.-A., Athanasiu, L., Selnes, P., Saltvedt, I., Sando, S.B., Ulstein, I., Djurovic, S., Fladby, T., Aarsland, D., Selbk, G., Ripke, S., Stefansson, K., Andreassen, O.A., Posthuma, D.: A genome-wide association study with 1,126,563 individuals identifies new risk loci for alzheimer's disease. Nature genetics **53**, 1276–1282 (2021)

[4] Baloni, P., Funk, C.C., Yan, J., Yurkovich, J.T., Kueider-Paisley, A., Nho, K., Heinken, A., Jia, W., Mahmoudiandehkordi, S., Louie, G., Saykin, A.J., Arnold, M., Kastenmller, G., Griffiths, W.J., Thiele, I., Kaddurah-Daouk, R., Price, N.D.: Metabolic network analysis reveals altered bile acid synthesis and metabolism in alzheimer's disease. Cell reports. Medicine **1**, 100138 (2020) https://doi.org/10.1016/j.xcrm.2020.100138

[5] Hurley, M.J., Bates, R., Macnaughtan, J., Schapira, A.H.V.: Bile acids and neurological disease. Pharmacology Therapeutics **240**, 108311 (2022) https://doi.org/10.1016/j.pharmthera.2022.108311

[6] Song, H., Liu, J., Wang, L., Hu, X., Li, J., Zhu, L., Pang, R., Zhang, A.: Tauroursodeoxycholic acid: a bile acid that may be used for the prevention and treatment of alzheimers disease. Frontiers in Neuroscience **18** (2024) https://doi.org/10.3389/fnins.2024.1348844

[7] Varma, V.R., Wang, Y., An, Y., Varma, S., Bilgel, M., Doshi, J., Legido-Quigley, C., Delgado, J.C., Oommen, A.M., Roberts, J.A., Wong, D.F., Davatzikos, C.,

Resnick, S.M., Troncoso, J.C., Pletnikova, O., OBrien, R., Hak, E., Baak, B.N., Pfeiffer, R., Baloni, P., Mohmoudiandehkordi, S., Nho, K., Kaddurah-Daouk, R., Bennett, D.A., Gadalla, S.M., Thambisetty, M.: Bile acid synthesis, modulation, and dementia: A metabolomic, transcriptomic, and pharmacoepidemiologic study. PLOS Medicine **18**(5), 1003615 (2021) https://doi.org/10.1371/journal.pmed.1003615

[18] Jansen, I.E., Savage, J.E., Watanabe, K., Bryois, J., Williams, D.M., Steinberg, S., Sealock, J., Karlsson, I.K., Hgg, S., Athanasiu, L., Voyle, N., Proitsi, P., Witoelar, A., Stringer, S., Aarsland, D., Almdahl, I.S., Andersen, F., Bergh, S., Bettella, F., Bjornsson, S., Brkhus, A., Brthen, G., Leeuw, C., Desikan, R.S., Djurovic, S., Dumitrescu, L., Fladby, T., Hohman, T.J., Jonsson, P.V., Kiddle, S.J., Rongve, A., Saltvedt, I., Sando, S.B., Selbk, G., Shoai, M., Skene, N.G., Snaedal, J., Stordal, E., Ulstein, I.D., Wang, Y., White, L.R., Hardy, J., Hjerling-Leffler, J., Sullivan, P.F., Flier, W.M., Dobson, R., Davis, L.K., Stefansson, H., Stefansson, K., Pedersen, N.L., Ripke, S., Andreassen, O.A., Posthuma, D.: Genome-wide meta-analysis identifies new loci and functional pathways influencing alzheimer's disease risk. Nature genetics **51**(30617256), 404–413 (2019) https://doi.org/10.1038/s41588-018-0311-9

[19] Kunkle, B.W., Grenier-Boley, B., Sims, R., Bis, J.C., Damotte, V., Naj, A.C., Boland, A., Vronskaya, M., Lee, S.J., Amlie-Wolf, A., Bellenguez, C., Frizatti, A., Chouraki, V., Martin, E.R., Sleegers, K., Badarinarayan, N., Jakobsdottir, J., Hamilton-Nelson, K.L., Moreno-Grau, S., Olaso, R., Raybould, R., Chen, Y., Kuzma, A.B., Hiltunen, M., Morgan, T., Ahmad, S., Vardarajan, B.N., Epelbaum, J., Hoffmann, P., Boada, M., Beecham, G.W., Garnier, J.-G., Harold, D., Fitzpatrick, A.L., Valladares, O., Moutet, M.-L., Gerrish, A., Smith, A.V., Qu, L., Bacq, D., Denning, N., Jian, X., Zhao, Y., Del Zompo, M., Fox, N.C., Choi, S.-H., Mateo, I., Hughes, J.T., Adams, H.H., Malamon, J., Sanchez-Garcia, F., Patel, Y., Brody, J.A., Dombroski, B.A., Naranjo, M.C.D., Daniilidou, M., Eiriksdottir, G., Mukherjee, S., Wallon, D., Uphill, J., Aspelund, T., Cantwell, L.B., Garzia, F., Galimberti, D., Hofer, E., Butkiewicz, M., Fin, B., Scarpini, E., Sarnowski, C., Bush, W.S., Meslage, S., Kornhuber, J., White, C.C., Song, Y., Barber, R.C., Engelborghs, S., Sordon, S., Voijnovic, D., Adams, P.M., Vandenberghe, R., Mayhaus, M., Cupples, L.A., Albert, M.S., De Deyn, P.P., Gu, W., Himali, J.J., Beekly, D., Squassina, A., Hartmann, A.M., Orellana, A., Blacker, D., Rodriguez-Rodriguez, E., Lovestone, S., Garcia, M.E., Doody, R.S., Munoz-Fernadez, C., Sussams, R., Lin, H., Fairchild, T.J., Benito, Y.A., Holmes, C., Karamuji-omi, H., Frosch, M.P., Thonberg, H., Maier, W., Roshchupkin, G., Ghetti, B., Giedraitis, V., Kawalia, A., Li, S., Huebinger, R.M., Kilander, L., Moebus, S., Hernndez, I., Kamboh, M.I., Brundin, R., Turton, J., Yang, Q., Katz, M.J., Concari, L., Lord, J., Beiser, A.S., Keene, C.D., Helisalmi, S., Kloszewska, I., Kukull, W.A., Koivisto, A.M., Lynch, A., Tarraga, L., Larson, E.B., Haapasalo, A., Lawlor, B., Mosley, T.H., Lipton, R.B., Solfrizzi, V., Gill, M., Longstreth, J. W T, Montine,

T.J., Frisardi, V., Diez-Fairen, M., Rivadeneira, F., Petersen, R.C., Deramecourt, V., Alvarez, I., Salani, F., Ciaramella, A., Boerwinkle, E., Reiman, E.M., Fievet, N., Rotter, J.I., Reisch, J.S., Hanon, O., Cupidi, C., Andre Uitterlinden, A.G., Royall, D.R., Dufouil, C., Maletta, R.G., Rojas, I., Sano, M., Brice, A., Cecchetti, R., George-Hyslop, P.S., Ritchie, K., Tsolaki, M., Tsuang, D.W., Dubois, B., Craig, D., Wu, C.-K., Soininen, H., Avramidou, D., Albin, R.L., Fratiglioni, L., Germanou, A., Apostolova, L.G., Keller, L., Koutroumani, M., Arnold, S.E., Panza, F., Gkatzima, O., Asthana, S., Hannequin, D., Whitehead, P., Atwood, C.S., Caffarra, P., Hampel, H., Quintela, I., Carracedo, ., Lannfelt, L., Rubinsztein, D.C., Barnes, L.L., Pasquier, F., Frlich, L., Barral, S., McGuinness, B., Beach, T.G., Johnston, J.A., Becker, J.T., Passmore, P., Bigio, E.H., Schott, J.M., Bird, T.D., Warren, J.D., Boeve, B.F., Lupton, M.K., Bowen, J.D., Proitsi, P., Boxer, A., Powell, J.F., Burke, J.R., Kauwe, J.S.K., Burns, J.M., Mancuso, M., Buxbaum, J.D., Bonuccelli, U., Cairns, N.J., McQuillin, A., Cao, C., Livingston, G., Carlson, C.S., Bass, N.J., Carlsson, C.M., Hardy, J., Carney, R.M., Bras, J., Carrasquillo, M.M., Guerreiro, R., Allen, M., Chui, H.C., Fisher, E., Masullo, C., Crocco, E.A., DeCarli, C., Bisceglio, G., Dick, M., Ma, L., Duara, R., Graff-Radford, N.R., Evans, D.A., Hodges, A., Faber, K.M., Scherer, M., Fallon, K.B., Riemenschneider, M., Fardo, D.W., Heun, R., Farlow, M.R., Klsch, H., Ferris, S., Leber, M., Foroud, T.M., Heuser, I., Galasko, D.R., Giegling, I., Gearing, M., Hll, M., Geschwind, D.H., Gilbert, J.R., Morris, J., Green, R.C., Mayo, K., Growdon, J.H., Feulner, T., Hamilton, R.L., Harrell, L.E., Drichel, D., Honig, L.S., Cushion, T.D., Huentelman, M.J., Hollingworth, P., Hulette, C.M., Hyman, B.T., Marshall, R., Jarvik, G.P., Meggy, A., Abner, E., Menzies, G.E., Jin, L.-W., Leonenko, G., Real, L.M., Jun, G.R., Baldwin, C.T., Grozeva, D., Karydas, A., Russo, G., Kaye, J.A., Kim, R., Jessen, F., Kowall, N.W., Vellas, B., Kramer, J.H., Vardy, E., LaFerla, F.M., Jckel, K.-H., Lah, J.J., Dichgans, M., Leverenz, J.B., Mann, D., Levey, A.I., Pickering-Brown, S., Lieberman, A.P., Klopp, N., Lunetta, K.L., Wichmann, H.-E., Lyketsos, C.G., Morgan, K., Marson, D.C., Brown, K., Martiniuk, F., Medway, C., Mash, D.C., Nthen, M.M., Masliah, E., Hooper, N.M., McCormick, W.C., Daniele, A., McCurry, S.M., Bayer, A., McDavid, A.N., Gallacher, J., McKee, A.C., Bussche, H., Mesulam, M., Brayne, C., Miller, B.L., Riedel-Heller, S., Miller, C.A., Miller, J.W., Al-Chalabi, A., Morris, J.C., Shaw, C.E., Myers, A.J., Wiltfang, J., O'Bryant, S., Olichney, J.M., Alvarez, V., Parisi, J.E., Singleton, A.B., Paulson, H.L., Collinge, J., Perry, W.R., Mead, S., Peskind, E., Cribbs, D.H., Rossor, M., Pierce, A., Ryan, N.S., Poon, W.W., Nacmias, B., Potter, H., Sorbi, S., Quinn, J.F., Sacchinelli, E., Raj, A., Spalletta, G., Raskind, M., Caltagirone, C., Boss, P., Orfei, M.D., Reisberg, B., Clarke, R., Reitz, C., Smith, A.D., Ringman, J.M., Warden, D., Roberson, E.D., Wilcock, G., Rogaeva, E., Bruni, A.C., Rosen, H.J., Gallo, M., Rosenberg, R.N., Ben-Shlomo, Y., Sager, M.A., Mecocci, P., Saykin, A.J., Pastor, P., Cuccaro, M.L., Vance, J.M., Schneider, J.A., Schneider, L.S., Slifer, S., Seeley, W.W., Smith, A.G., Sonnen, J.A., Spina, S., Stern, R.A., Swerdlow, R.H., Tang, M., Tanzi, R.E., Trojanowski, J.Q., Troncoso, J.C., Van Deerlin, V.M., Van Eldik,

L.J., Vinters, H.V., Vonsattel, J.P., Weintraub, S., Welsh-Bohmer, K.A., Wilhelmsen, K.C., Williamson, J., Wingo, T.S., Woltjer, R.L., Wright, C.B., Yu, C.-E., Yu, L., Saba, Y., Pilotto, A., Bullido, M.J., Peters, O., Crane, P.K., Bennett, D., Bosco, P., Coto, E., Boccardi, V., De Jager, P.L., Lleo, A., Warner, N., Lopez, O.L., Ingelsson, M., Deloukas, P., Cruchaga, C., Graff, C., Gwilliam, R., Fornage, M., Goate, A.M., Sanchez-Juan, P., Kehoe, P.G., Amin, N., Ertekin-Taner, N., Berr, C., Debette, S., Love, S., Launer, L.J., Younkin, S.G., Dartigues, J.-F., Corcoran, C., Ikram, M.A., Dickson, D.W., Nicolas, G., Campion, D., Tschanz, J., Schmidt, H., Hakonarson, H., Clarimon, J., Munger, R., Schmidt, R., Farrer, L.A., Van Broeckhoven, C., C O'Donovan, M., DeStefano, A.L., Jones, L., Haines, J.L., Deleuze, J.-F., Owen, M.J., Gudnason, V., Mayeux, R., Escott-Price, V., Psaty, B.M., Ramirez, A., Wang, L.-S., Ruiz, A., Duijn, C.M., Holmans, P.A., Seshadri, S., Williams, J., Amouyel, P., Schellenberg, G.D., Lambert, J.-C., Pericak-Vance, M.A., (ADGC), A.D.G.C., (EADI), E.A.D.I., Heart, C., Genomic Epidemiology Consortium (CHARGE), A.R., Genetic, AD/Defining Genetic, P., Alzheimers Disease Consortium (GERAD/PERADES), E.R.: Genetic meta-analysis of diagnosed alzheimer's disease identifies new risk loci and implicates a$\beta$, tau, immunity and lipid processing. Nature genetics **51**(30820047), 414–430 (2019) https://doi.org/10.1038/s41588-019-0358-2

[20] Wingo, A.P., Liu, Y., Gerasimov, E.S., Gockley, J., Logsdon, B.A., Duong, D.M., Dammer, E.B., Robins, C., Beach, T.G., Reiman, E.M., Epstein, M.P., De Jager, P.L., Lah, J.J., Bennett, D.A., Seyfried, N.T., Levey, A.I., Wingo, T.S.: Integrating human brain proteomes with genome-wide association data implicates new proteins in alzheimer's disease pathogenesis. Nature genetics **53**(33510477), 143–146 (2021) https://doi.org/10.1038/s41588-020-00773-z

[21] Taubes, A., Nova, P., Zalocusky, K.A., Kosti, I., Bicak, M., Zilberter, M.Y., Hao, Y., Yoon, S.Y., Oskotsky, T., Pineda, S., Chen, B., Aery Jones, E.A., Choudhary, K., Grone, B., Balestra, M.E., Chaudhry, F., Paranjpe, I., De Freitas, J., Koutsodendris, N., Chen, N., Wang, C., Chang, W., An, A., Glicksberg, B.S., Sirota, M., Huang, Y.: Experimental and real-world evidence supporting the computational repurposing of bumetanide for apoe4-related alzheimers disease. Nature Aging **1**(10), 932–947 (2021) https://doi.org/10.1038/s43587-021-00122-7

[22] Lin, Y.-T., Seo, J., Gao, F., Feldman, H.M., Wen, H.-L., Penney, J., Cam, H.P., Gjoneska, E., Raja, W.K., Cheng, J., Rueda, R., Kritskiy, O., Abdurrob, F., Peng, Z., Milo, B., Yu, C.J., Elmsaouri, S., Dey, D., Ko, T., Yankner, B.A., Tsai, L.-H.: Apoe4 causes widespread molecular and cellular alterations associated with alzheimer's disease phenotypes in human ipsc-derived brain cell types. Neuron **98**(29861287), 1141–11547 (2018) https://doi.org/10.1016/j.neuron.2018.05.008

[23] Cicchetti, F., David, L.S., Siddu, A., Denis, H.L.: Cysteamine as a novel disease-modifying compound for parkinson's disease: Over a decade of research supporting a clinical trial. Neurobiology of Disease **130**, 104530 (2019) https://doi.org/10.1016/j.nbd.2019.104530
